# Theoretical consequences of the Mutagenic Chain Reaction for manipulating natural populations

**DOI:** 10.1101/018986

**Authors:** Robert L. Unckless, Philipp W. Messer, Andrew G. Clark

## Abstract

The use of recombinant genetic technologies for population manipulation has mostly remained an abstract idea due to the lack of a suitable means to drive novel gene constructs to high frequency in populations. Recently Gantz and Bier showed that the use of CRISPR/Cas9 technology could provide an artificial drive mechanism, the so-called Mutagenic Chain Reaction (MCR), which could lead to rapid fixation of even a deleterious introduced allele. We establish the equivalence of this system to models of meiotic drive and review the results of simple models showing that, when there is a fitness cost to the MCR allele, an internal equilibrium exists that is usually unstable. Introductions must be at a frequency above this critical point for the successful invasion of the MCR allele. These modeling results have important implications for application of MCR in natural populations.

## Introduction

The prospect of introducing a novel gene into a population and having it spread to high frequency holds great promise for biological control. Such technologies could improve crop yields by pest reduction, stop the spread of vector-borne disease, and potentially alleviate environmental problems (Bourtzis and Hendrichs 2014). But the possibility of uncontrolled spread of this artificial genetic material once introduced is a cause of significant concern (Bohannon 2015). With few exceptions (Hoffmann *et al.* 2011), the practicality of such introductions has been limited by the lack of a means to ensure the spread of the genetic material of interest through a population. In a recent paper, Gantz and Bier (Gantz and Bier 2015) describe the Mutagenic Chain Reaction (MCR), an approach that employs the CRISPR/Cas9 system to drive a mutation to high frequency in a population, making gene replacement at the population level practical for any species that can be made to accept a transgene in the laboratory.

We envision three classes of introductions. First, the introduced genetic material could be beneficial to the organism. Imagine a mutation that confers resistance to malaria in *Anopheles* mosquitoes or allows an endangered species to tolerate a new environmental stress. This is analogous to the use of the virus suppressing *Wolbachia* to combat Dengue Fever in *Aedes* mosquitoes (Bull and Turelli 2013). Second, the mutation could be neutral in a standard environment. In this case, a mutation that confers insecticide susceptibility could be introduced into an agricultural pest population without the use of that particular pesticide, then when the mutation is near fixation, the pesticide could be applied, thereby reducing the population severely. Finally, a mutation might be deleterious regardless of environment. This last scenario is similar to sterile male techniques that have been discussed for decades (Foster *et al.* 1972; Prout 1978; Bourtzis and Hendrichs 2014). For example, a life-shortening mutation in mosquitoes that does not allow for a complete incubation period for Dengue virus could severely reduce transmission rates. In this case, the rate of conversion via the MCR process must outweigh the fitness cost to the organism. Here we model all three scenarios, find an internal equilibrium for those mutations with fitness costs, and interpret these results in terms of practical applications.

## Modeling MCR

The essential feature of MCR with respect to population frequency dynamics is that heterozygotes produce an excess of gametes bearing the MCR construct. Meiotic drive and gene conversion are two quite distinct mechanisms that also produce such non-Mendelian gametic counts. The population genetic consequences of meiotic drive (Prout 1953) and of gene conversion (Gutz and Leslie 1976) have both been shown to result in rapid fixation. When balanced by opposing natural selection, both mechanisms can produce internal equilibria (Hiraizumi *et al.* 1960; Walsh 1983). In fact, the model for which MCR is a special case is given by Hartl (Hartl 1970), and here we reparameterize this model to make clear how it relates to the biological process of MCR and its relevance to biological control. First, we posit that the MCR allele will have a fitness cost in homozygous individuals that we designate as *s*, and the dominance term *h* represents the dominance of the MCR allele fitness costs in heterozygotes. Thus, the wildtype homozygote fitness is 1.0, the heterozygote for the MCR allele has fitness 1-*hs*, and the homozygote for the MCR allele has fitness 1- *s*. While most of the interesting modeling occurs when *s* is positive (the allele is costly), our model does allow for the introduction of a beneficial (*s <* 0) or neutral (*s* = 0) allele. We let *c* represents the rate of CRISPR-mediated conversion from wildtype to the MCR allele in heterozygotes. Finally 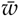 is the mean fitness that normalizes the recursion so that allele frequencies sum to one. Considering all these processes, we can write the recursion that gives the frequency of the MCR allele in the following generation, given its current frequency *q*:

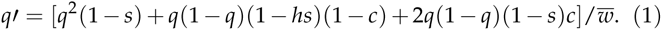

This model assumes a large, random mating, panmictic population and that CRISPR-mediated gene conversions happens in heterozygote embryos.

To get some sense of the behavior of this model, we show some plots of the frequency of the MCR construct, starting at an initial frequency of 0.001, and a range of selection coefficients and conversion rates (Figure 1). Note that the rate of spread is impressively rapid, even in cases where the MCR construct confers a 25 percent reduction in fitness, and with a conversion efficiency of only 80 percent. These simulation results certainly underscore the potential hazard of allowing MCR to run uncontrolled in natural populations.

**Figure 1.**
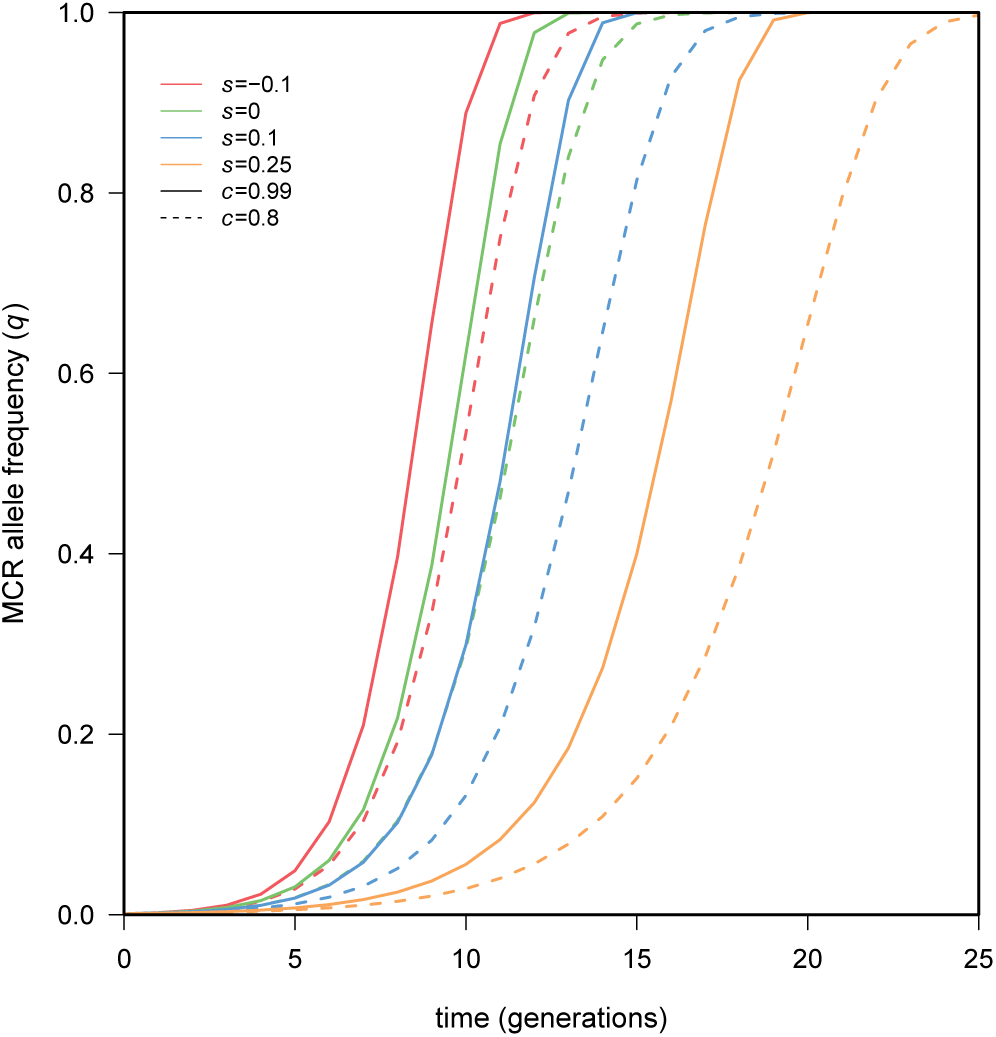
Trajectories of introduced MCR alleles reveal that even deleterious alleles sweep to fixation very quickly. Only parameter sets leading to fixation are presented, and all cases shown assume that fitness costs are recessive.

This model can admit a single internal equilibrium:

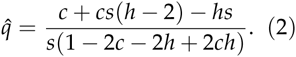

The stability of this internal equilibrium depends on the values of the three parameters of the model (Figures 2,3, Appendix 1). Briefly, if fitness costs of the MCR allele are dominant (*h* = 1), then the internal equilibrium is unstable. For MCR to successfully invade, there is a constraint on the fitness cost 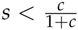. If fitness costs are recessive, the internal equilibrium is stable if *c <* 1/2 and unstable if *c >* 1/2. For additive fitness, the internal equilibrium is always unstable (Figure 3).

**Figure 2.**
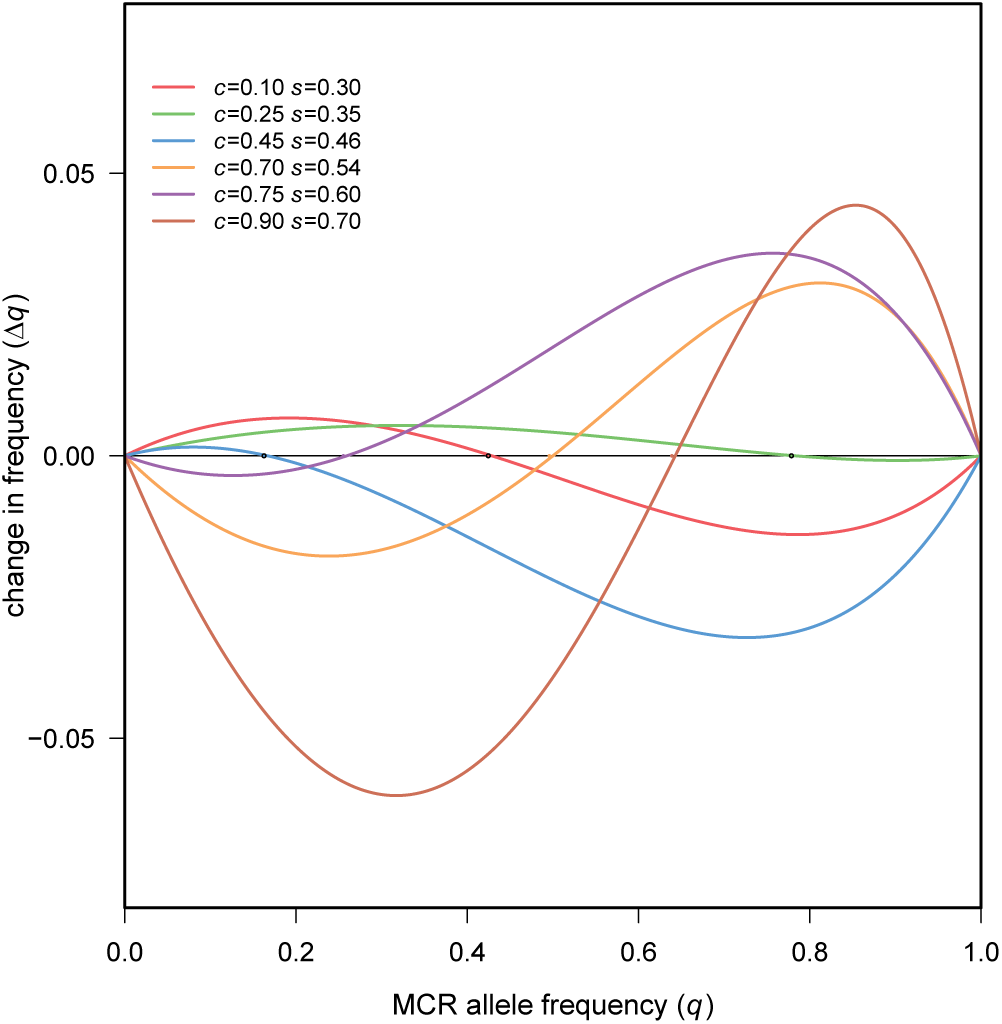
The change in MCR allele frequency (Δ*q*) near equilibrium reveals the local stability of the equilibrium. Positive slopes are associated with unstable equilibria and negative slopes are associated with stable equilibria (black borders on circles). Fitness costs are recessive (*h* = 0).

**Figure 3.**
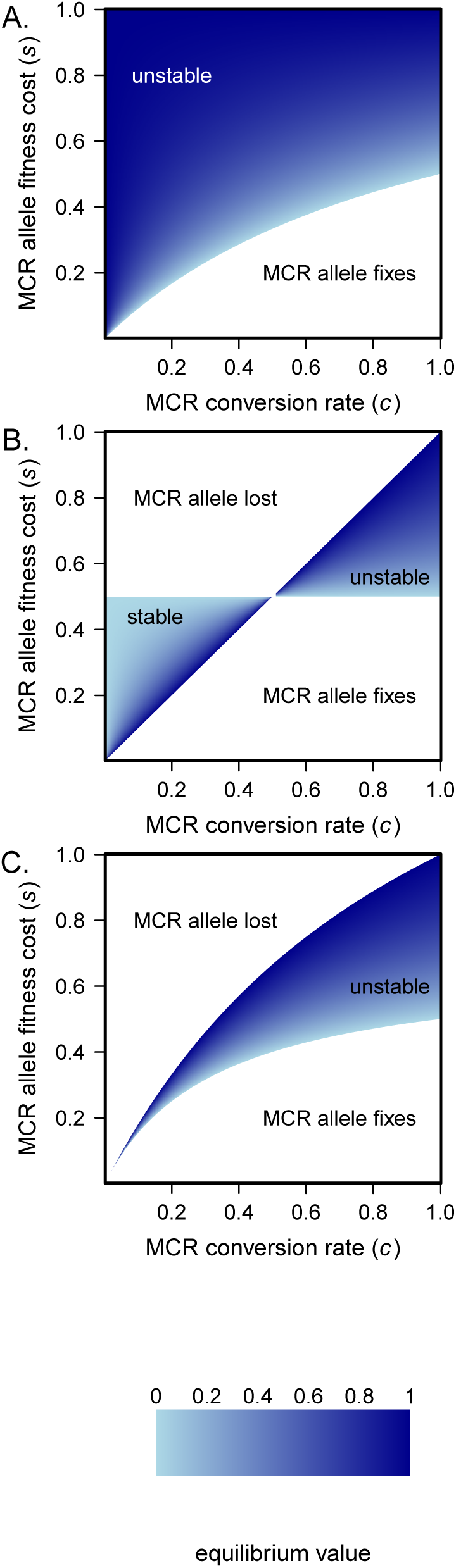
Equilibria when fitness costs are A) dominant (*h* = 1), B) recessive (*h* = 0) and C) additive (*h* = 1/2). Parameter space leading to fixation, loss, stable or unstable equilibrium of the MCR allele are noted on the figures.

In the case of biological control applications, the goal will be to attain fixation of the MCR allele, and so the construct must be introduced into the wild population at sufficient frequency to exceed the unstable equilibrium frequency. This is reminiscent of the spread of *Wolbachia* through populations (Turelli and Hoffmann 1995). In both cases, insufficient initial frequency will result in loss from the population. From the point of view of biosafety, we would want any escaped version of the MCR to have an elevated unstable equilibrium, so that the population would only be able to exceed this threshold with a concerted effort by researchers consciously working toward that goal. Figure 3 shows the location of this critical equilibrium frequency, and shows that the equilibrium frequency is elevated when fitness costs are high, when dominance is reduced, and when conversion rates to the MCR allele relatively low. One possible route to increase the safety of an introduction would be to engineer means to assure low conversion frequency and high fitness cost, at the same time yielding an equilibrium frequency that is practical in terms of ability to produce and release the engineered organisms. Of course, one should expect that all parameters in the system would be free to evolve once released into a natural population, necessitating other safeguards (see Discussion).

Of particular interest is the likelihood that an MCR allele would escape stochastic loss and sweep to fixation either through purposeful or accidental introduction. This scenario might be most relevant when considering invasion of the MCR construct into a non-target species, where initial frequency may be quite low. This type of modeling requires the inclusion of genetic drift and models are therefore stochastic. We restrict our analysis to cases parameter space where, in a deterministic model, fixation occurs and there is no internal equilibrium (see above). Again, one would not engineer the system to have this property, but an MCR that escapes into a non-target species might. It follows that in most of the parameter space, the fate of the MCR allele is determined when it is quite rare and we can ignore terms involving *q*^2^. Equation (1) then becomes

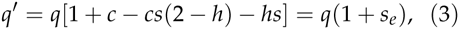

resembling standard exponential growth with an effective selection coefficient *s*_*e*_= *hs*(*c –* 1) + (1 – 2*s*)*c*. In this case, the probability of fixation of the MCR allele can be approximated as

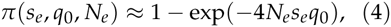

where *N*_*e*_ is the variance effective population size and *q*_0_ is the starting frequency of the MCR allele. Furthermore, the minimum frequency at which the MCR allele needs to be introduced into the population to escape stochastic loss is *q*_critical_ *>* 1/(2*N*_*e*_ *s*_*e*_). This means that in the Wright-Fisher scenario, the MCR allele must be introduced in 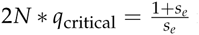 individuals to be relatively certain that it will escape stochastic loss. Figure 4 shows that the approximation in Equation (4) provides a reasonable match to forward simulations but slightly overestimates fixation probability especially as *c* becomes large and *s* becomes small.

**Figure 4.**
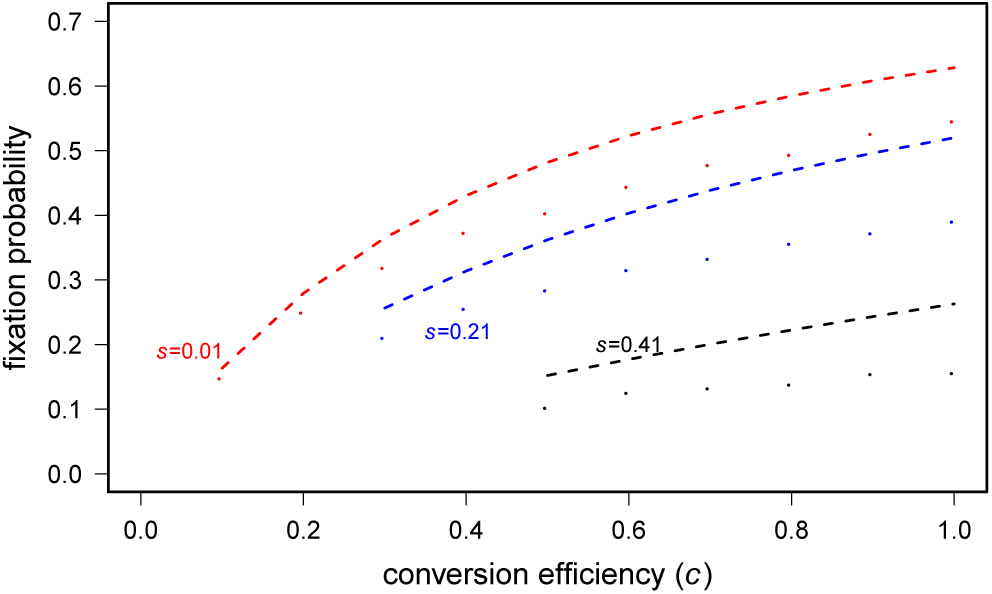
Probability of fixation when fitness costs are recessive (*h* = 0). Only cases leading to fixation without equilibrium are plotted. All simulations assume a population size of 10,000, a single copy of the MCR allele to start. Points represent the proportion of simulation realizations with MCR allele fixation. Lines represent analytical approximations, using the scaling *N*_*e*_ = *N*/(1 + *s*_*e*_) for the Wright-Fisher model (Uecker and Hermisson 2011).

Our approximation for the effective selection coefficient also allows for an estimate of the time it takes for the allele to become prevalent in the population. For example, assuming exponential growth at rate *s*_*e*_, the MCR allele is expected to have invaded half the population after approximately

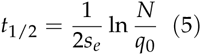

generations.

## Discussion

Several factors not modeled here could alter the spread of an MCR allele. First, the MCR allele could experience a mutation rendering it unable, by itself, to convert the homolog to the MCR state. This could severely limit the spread of the MCR allele when it carries fitness costs (*s >* 0). In fact, this could even be used as a method for controlling the MCR allele. With three alleles: wildtype, active MCR, and broken MCR, we imagine that the active MCR converts the wildtype but cannot reconvert the broken MCR allele, and that the broken MCR allele has higher fitness than the active MCR allele (it could be engineered that way). In this case, the introduction of the broken MCR allele should spread since it is both resistant to conversion from the active MCR allele and will have higher fitness than the MCR allele as well.

Another possibility is that genetic variation for conversion efficiency may segregate in populations. This could slow the spread of the MCR allele or even change the invasion dynamics - if some backgrounds render *c <* 1/2, the equilibrium could become stable. Population geneticists have modeled this situation with a second locus that serves to modify the strength of meiotic drive at the driven locus (Prout *et al.* 1973). The conclusion from this analysis is that modifier loci can easily select alleles that impact the population dynamics, and when they occur in stable equilibria, they generally are in linkage disequilibrium with the driven locus. The very rapid fixation of MCR may preclude the invasion of such modifiers, but in any case there is already machinery for analysis of such modifiers.

Finally, our employment of the Wright-Fisher model surely overestimates the variance effective population size of most real populations. Due to population structure, mating system, or several other factors, effective population sizes are likely much less than census population sizes (Palstra and Ruzzante 2008). In fact, in many insect, such as mosquitoes, measured effective population sizes are on the order of tens of thousands, several orders of magnitude less than the true census population size (Athrey *et al.* 2012).

These results show that the MCR may provide an effective means for population replacement, and that the speed of the process presents reason for considerable caution before considering a field release of such a construct. In fact, there are conditions in which accidental introductions of a single individual can lead to fixation of the MCR allele even with significant fitness consequences to the individual. For example, an allele with perfect conversion efficiency and a selective cost of 0.41 escapes stochastic loss and sweeps to fixation nearly 20 percent of the time. Thorough quantitative modeling of MCR population dynamics is strongly warranted, not only to put bounds on the frequency trajectories expected from release of an MCR, but also possible choke points for controlling and preventing the expansion of an escaped or mutated MCR allele in a natural population.

Several additional lines of inquiry will be the subject of future work. For example, in a structured population a balance between conversion and migration might facilitate the spread of the MCR allele. Also, one potential application of MCR is to purposefully drive pest populations to extinction. In this case it would be useful to predict the expected time to extinction and the likelihood that the population can rescue itself before extinction. A population might be rescued by an allele in the standing genetic variation or new mutation that is resistant to conversion (Orr and Unckless 2014).

## Appendix 1

To determine the stability of the equilibrium in Equation (2), we first find the change in MCR allele frequency in a single generation:

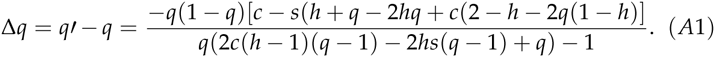

At equilibrium (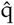, Eq. 2), the eigenvalue for equation A1 is

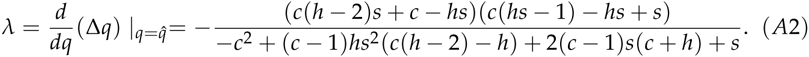

For an MCR allele with dominant fitness costs (*h* = 1), the equilibrium in Equation (2) is between zero and one only if 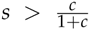, otherwise the MCR allele always fixes. However, when 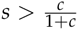, the eigenvalue in equation (A2) is positive, and therefore the equilibrium is unstable. This also means that MCR alleles with a dominant fitness cost greater than *s* = 0.5 cannot successfully invade, since no interior equilibrium exists, and the equilibrium at *q* = 1 is unstable.

If the MCR allele carries recessive fitness costs (*h* = 0), the equilibrium in Equation (2) is between zero and one, if one of two sets of conditions are met. First, if *c <* 1/2, the following must be met: *c < s <* 1/2. Alternatively, if *c >* 1/2, the following must be met: 1/2 *< s < c*. The internal equilibrium is stable when *c <* 1/2, because for the eigenvalue in Equation (2) to be positive, either *s >* 1/2 or *c > s*, both conditions would not permit an internal equilibrium. If *c >* 1/2, then the internal equilibrium is always unstable.

Finally, in the case of additive fitness (*h* = 1/2), an internal equilibrium exists if 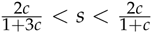, and this equilibrium is always unstable. The general case, where *h* is a free parameter between zero and one is solvable but complex and therefore not presented here.

